# Castor is a temporal transcription factor that specifies early born central complex neuron identity

**DOI:** 10.1101/2024.08.22.609207

**Authors:** Noah R. Dillon, Chris Q. Doe

**Affiliations:** Institute of Neuroscience, Howard Hughes Medical Institute, University of Oregon, Eugene, OR 97403

**Keywords:** Castor, temporal specification, neuroblast, central complex, *Drosophila*

## Abstract

The generation of neuronal diversity is important for brain function, but how diversity is generated is incompletely understood. We used the development of the *Drosophila* central complex (CX) to address this question. The CX develops from eight bilateral Type 2 neuroblasts (T2NBs), which generate hundreds of different neuronal types. T2NBs express broad opposing temporal gradients of RNA-binding proteins. It remains unknown whether these protein gradients are sufficient to directly generate all known neuronal diversity, or whether there are temporal transcription factors (TTFs) with narrow expression windows that each specify a small subset of CX neuron identities. Multiple candidate TTFs have been identified, but their function remains uncharacterized. Here, we show that: (i) the adult E-PG neurons are born from early larval T2NBs; (ii) the candidate TTF Castor is expressed transiently in early larval T2NBs when E-PG and P-EN neurons are born; and (iii) that Castor is required to specify early born E-PG and P-EN neuron identities. We conclude that Castor is a TTF in larval T2NB lineages that specifies multiple, early born CX neuron identities.

**Summary:** Castor acts in Type 2 neuroblast lineages as a temporal transcription factor to specify adult central complex columnar neuron identity.

## Introduction

Generating a complex brain requires neural stem cells to generate a large and diverse set of cell types. The neural stem cells in *Drosophila*, known as neuroblasts (NBs), generate neuronal diversity through a combination of spatial and temporal patterning mechanisms (Doe, 2017; El-Danaf et al., 2023). Similar results have been observed in mammals (Alsio et al., 2013; Clark et al., 2019; Elliott et al., 2008; Frith et al., 2024; Javed et al., 2020; Liu et al., 2020; Mattar and Cayouette, 2015; Mattar et al., 2021), highlighting the importance of temporal patterning in generating neuronal diversity across species. In *Drosophila*, these processes have been most extensively studied in embryonic NB lineages where narrow windows of temporal transcription factors (TTFs) specify distinct neuronal identities (Cleary and Doe, 2006; Grosskortenhaus et al., 2006; Isshiki et al., 2001; Meng et al., 2019; Meng et al., 2020; Moris-Sanz et al., 2015; Novotny et al., 2002; Pearson and Doe, 2003; Seroka and Doe, 2019; Tran and Doe, 2008). Whether larval NBs undergo similar TTF windows remains understudied.

The larval central brain contains ∼100 NB lineages per hemibrain (Pereanu and Hartenstein, 2006). There are two types of NBs based on their lineages: Type 1 NBs generate a series of ganglion mother cells that divide once to produce a pair of post-mitotic neurons, whereas Type 2 NBs (T2NBs) generate a series of intermediate neural progenitors (INPs) that each generate 4-6 ganglion mother cells that divide to produce a pair of post-mitotic neurons (Bello et al., 2008; Boone and Doe, 2008; Bowman et al., 2008). This T2NB division pattern is analogous to the outer subventricular zone lineages of the primate cortex (Holguera and Desplan, 2018; El-Danaf et al., 2023); thus, it is important to understand how these larval lineages, which closely resemble mammalian neural stem cells, are patterned to generate neuronal diversity. There are only eight T2NBs per hemibrain, with each lineage having a unique spatial identity that generates neurons with lineage-specific morphology (Andrade et al., 2019; Pereanu and Hartenstein, 2006; Riebli et al., 2013; Yang et al., 2013). Previous work has suggested that temporal patterning in both the T2NBs and INPs act in combination to generate neuronal diversity (Bayraktar and Doe, 2013).

Larval NBs express broad and opposing temporal gradients of two RNA-binding proteins: high levels of IGF-II mRNA-binding protein (Imp) in early stages and high levels of Syncrip in late stages (Fig. 1A) (Liu et al., 2015; Ren et al., 2017; Syed et al., 2017). The short burst of Seven-up (Svp) expression has been identified as a switching factor to initiate the temporal Imp-to-Syncrip transition in larval T2NBs (Fig. 1A) (Ren et al., 2017; Syed et al., 2017). We have previously shown that Svp acts in T2NBs to initiate a switch for transitioning the generation of early born fates to late born fates (Dillon et al., 2024); thus, temporally expressed factors in T2NBs are required for specifying neuronal identities. Previous work has demonstrated the importance of TTFs in the INPs to generate neuron subtype diversity (Bayraktar and Doe, 2013; Sullivan et al., 2019). Several candidate TTFs have been shown to have narrow temporal windows of expression in the T2NBs (Castor, Chinmo, Broad, and E93) (Ren et al., 2017; Syed et al., 2017) but whether these factors are required to specify different neuronal subtypes remains unknown.

**Figure 1.**
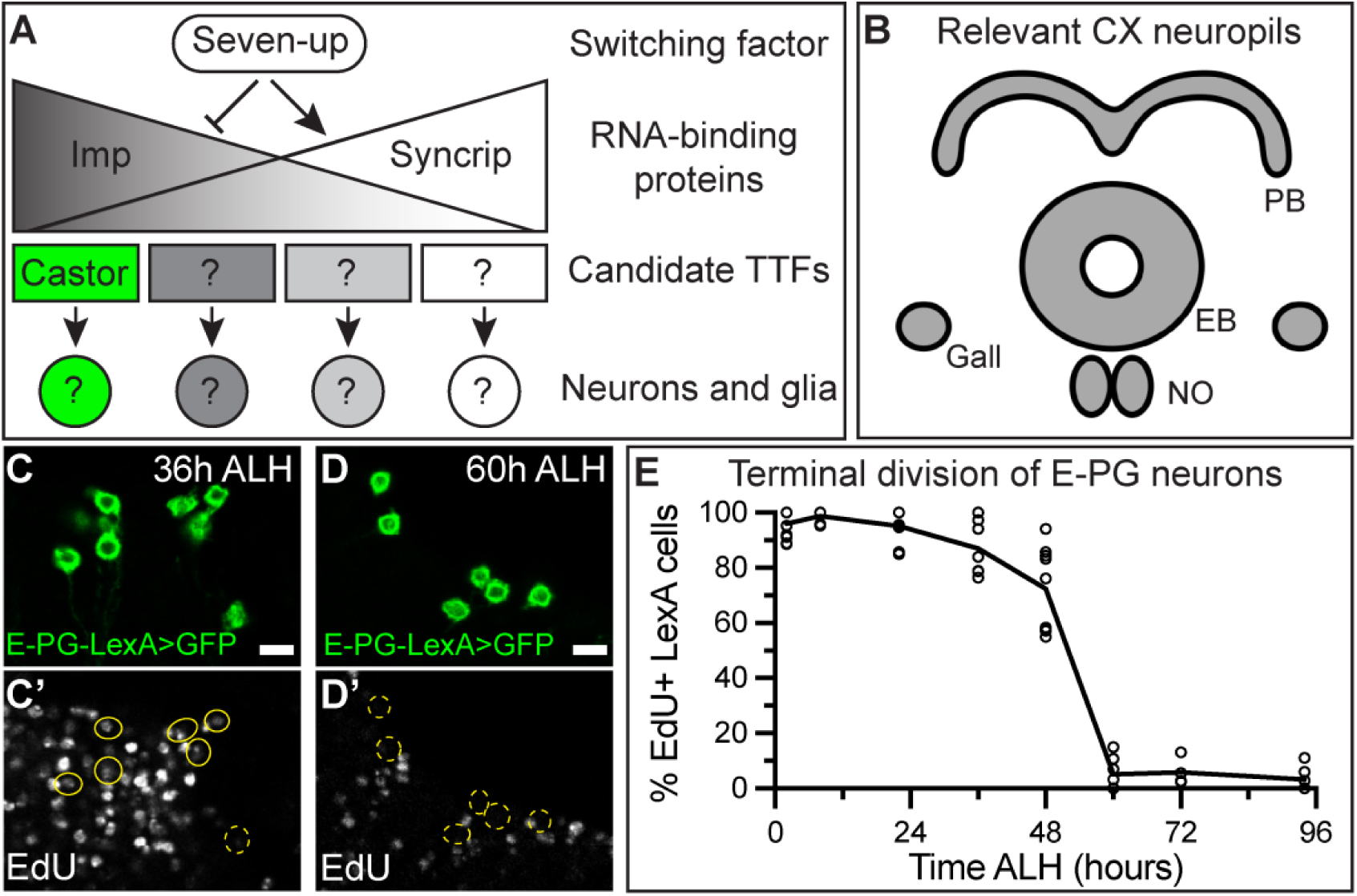
E-PG neurons are born early in Type 2 neuroblast lineages. (A) Larval neuroblasts express broad temporal gradients of the RNA-binding proteins Imp and Syncrip and shorter temporal windows of candidate temporal transcription factors (TTFs) such as Castor in the early larval T2NBs (Ren et al., 2017; Syed et al., 2017). (B) Schematic of central complex (CX) neuropils of the adult brain relevant to this study. PB, protocerebral bridge; EB, ellipsoid body; NO, noduli. (C-E) EdU labeling shows nearly all E-PG neurons are being generated prior to 36h ALH (EdU-positive) (C-C’) but are post-mitotic (EdU-negative) by 60h ALH (D-D’). Similar results are seen for P-EN neurons (Dillon et al., 2024). (E) Quantification. Each dot represents one adult brain. Lines connected at mean for each timepoint. For each timepoint, *n* = 4-11 brains. In all images, LexA+ neurons driving membrane-bound GFP are in green and outlined in yellow; solid line indicates positive for EdU, dashed line indicates negative for EdU. All scale bars: 5 μm.

T2NB lineages generate the majority of the adult *Drosophila* central complex (CX) with connectomes showing this region of the central brain to contain hundreds of morphologically distinct neuron subtypes (Franconville et al., 2018; Hulse et al., 2021). One group of CX neurons of interest have been the columnar neurons, which form neural circuits responsible for locomotion and spatial navigation (Giraldo et al., 2018; Green et al., 2017; Green et al., 2019; Turner-Evans et al., 2020). Columnar neurons are characterized by their axonal and dendritic projections into CX neuropils (Fig. 1B). For example, in this study we focused on E-PG neurons that send dendrites to the ellipsoid body (<u>E</u>B) and axons to the protocerebral bridge (<u>P</u>B) and <u>g</u>all; and P-EN neurons that send dendrites to the <u>P</u>B and axons to the <u>E</u>B and noduli (<u>N</u>O) (Wolff and Rubin, 2018; Wolff et al., 2015). Notably, the E-PG and P-EN neurons are within the same navigational circuit (Green et al., 2017; Green et al., 2019). Understanding the development of these neurons may shed light on how NB lineages generate complex behavioral circuits.

In this study, we show that Castor is a TTF in Type 2 (T2) lineages that specify early born CX neuronal identity. We find that E-PG neurons are born in an early T2NB window and have molecularly distinct markers that distinguish their identity from another group of early born neurons, the P-EN neurons. We find that Svp is required to restrict the generation of E-PG and P-EN neurons (this work and (Dillon et al., 2024)). We define the expression of the candidate TTF Castor in T2NBs to be transiently expressed in T2 lineages. We use T2NB lineage-specific manipulations of Castor during larval development to perform CRISPR/Cas9 knockouts of *castor* or ectopically extend the temporal expression of Castor in T2NBs to assay the impacts on adult CX neuron identities. We show that Castor is required to specify the early born E-PG and P-EN neuron identities. Additionally, we show Castor is sufficient to produce extra adult P-EN neurons but not E-PG neurons. This is the first study to identify a T2NB lineage TTF, Castor, expressed in a narrow T2NB temporal window that subdivides the previously known broad RNA-binding protein gradients, and to show it is necessary and sufficient to temporally specify columnar neurons in the adult CX.

## Results

### E-PG neurons are born early in Type 2 neuroblast lineages

We previously birth dated the columnar subtype of P-EN neurons to the early temporal window in the T2NB lineages (Dillon et al., 2024) and derived from young INPs (Sullivan et al., 2019). To identify additional early born neurons, we performed our 5-ethynyl-2′-deoxyuridine (EdU) drop out birth dating to find an additional early born columnar identity. We found that E-PG (R60D05-LexA) neurons become postmitotic, thus do not incorporate EdU, between 36 h and 60 h ALH (after larval hatching, Fig. 1C-E). We utilized cell cycle data for T2NBs, INPs and ganglion mother cells to determine the time from E-PG terminal division to birth from the parental T2NB (Bello et al., 2008; Bowman et al., 2008; Homem et al., 2013). E-PG neurons are derived from old INPs (Sullivan et al., 2019), which divide every 2 h for 4-6 divisions (Bello et al., 2008; Bowman et al., 2008) and are birthed from the parental T2NB 12 h before producing neurons (Homem et al., 2013); thus, neurons born from old INPs are derived from the parental T2NB 16-18 h prior to terminal division. We conclude that E-PG neurons are born from T2NBs between 20 h and 44 h ALH; this is similar to previous birth dating approaches for E-PG neurons (Sullivan et al., 2019) and is during the Castor expression window in T2NBs (Ren et al., 2017; Syed et al., 2017).

### E-PG and P-EN neurons have distinct molecular identities

Here, we focus on the E-PG and P-EN neurons as both are generated early from the same T2NBs but in different INP lineages (Dillon et al., 2024; Sullivan et al., 2019; Yang et al., 2013). Previous work has shown that P-EN neurons have a distinct molecular identity based on the expression of R12D09-LexA (Wolff et al., 2015) and the transcription factors Cut (Dillon et al., 2024; Epiney et al., 2023) and Runt (Sullivan et al., 2019). In contrast, E-PG neurons express the R60D05-LexA (Wolff et al., 2015) and the transcription factor Toy (Sullivan et al., 2019). To determine additional markers to distinguish E-PG neurons, we used a single-cell RNA-sequencing atlas of adult T2NB-derived neurons to identify markers that distinguish E-PG and P-EN neurons (Epiney et al., 2023). We found the transcription factor Dac was expressed in adult E-PG neurons (Fig. 2A-A’’, C) but not in adult P-EN neurons (Fig. 2B-B’’, C). We conclude that E-PG and P-EN neurons have multiple markers that distinguish their molecular identities (Fig. 2D). We use these markers in subsequent analyses as readouts of neuronal identities.

**Figure 2.**
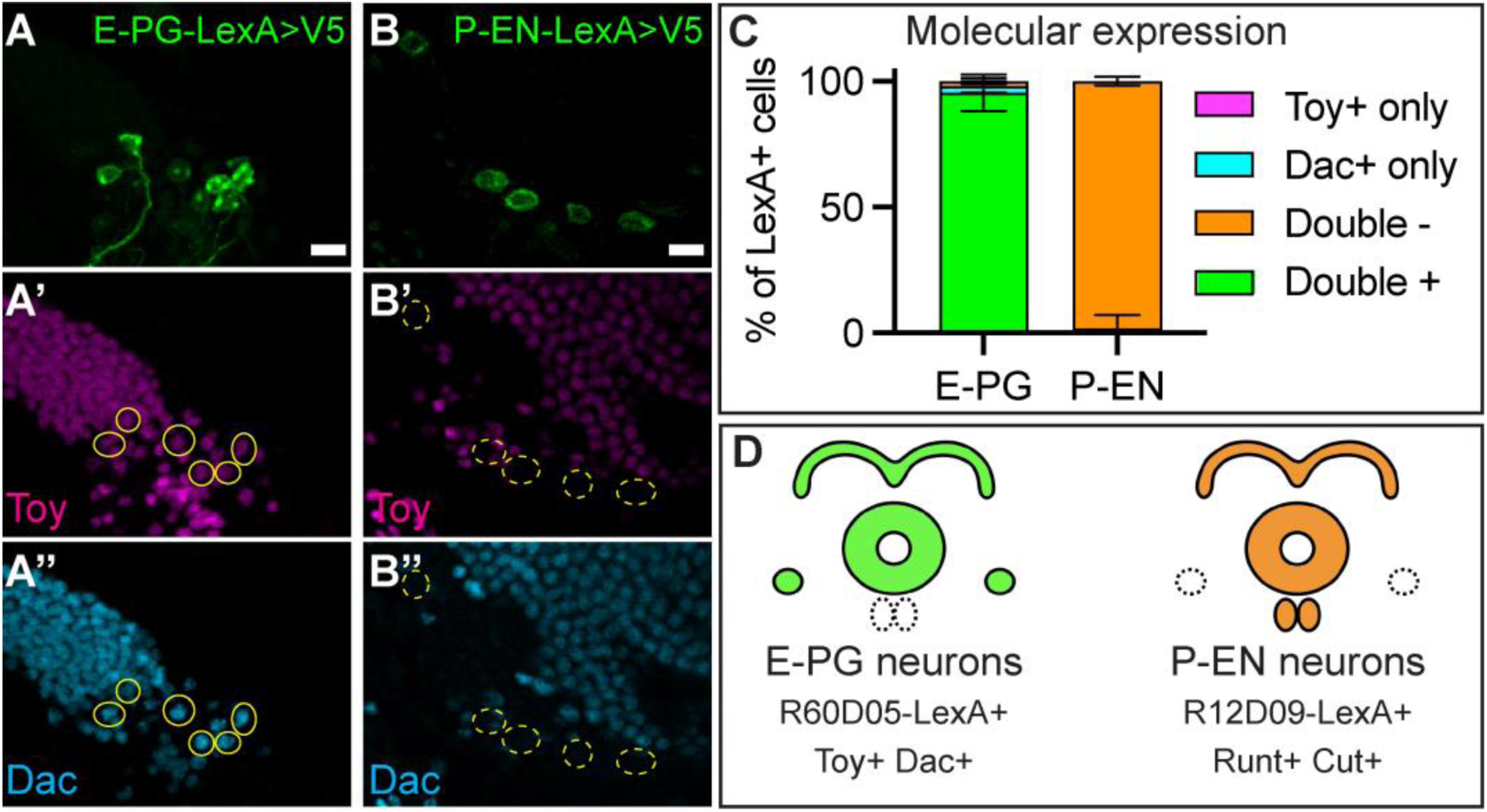
E-PG and P-EN neurons have distinct molecular identities. (A-C) Adult E-PG neurons express Toy and Dac (A-A’’) whereas adult P-EN neurons do not express Toy or Dac. (B-B’’). (C) Quantification. Error bars show standard error of the mean. For each genotype, *n* = 4-6 brains. (D) Summary. E-PG and P-EN are both early born from T2NBs but differ in neuropil targeting and molecular markers (Dillon et al., 2024). In all images, LexA+ neurons driving membrane-bound V5 are in green and outlined in yellow; solid line indicates double positive for markers, dashed line indicates double negative for markers; Toy, magenta; Dac, cyan. All scale bars: 5 μm.

### Seven-up is required to restrict E-PG production in early Type 2 neuroblast lineages

Previous work has shown that the loss of Svp in T2NB lineages leads to the production of extra early born neuron identities, such as P-EN neurons, by extending the early temporal window (Dillon et al., 2024; Ren et al., 2017). To confirm that early temporal factors are similarly required to specify E-PG neurons, we tested if loss of Svp in the T2NBs could produce extra E-PG neurons. We used previously validated, lineage specific CRISPR/Cas9 lines (Dillon et al., 2024) to generate Svp knockouts (Svp-KO) in the T2NBs with the T2NB driver Pnt-Gal4 (Zhu et al., 2011) in combination with the R60D05-LexA driver expressing a membrane-bound V5 epitope to label adult E-PGs neurons. We found that loss of Svp leads to an expansion of E-PG neuron numbers (Fig. 3A-B, E). Additionally, Svp-KOs led to defective EB morphology, with an incomplete closure of the EB neuropil (Fig. 3A-B; Fig. S1). The ectopic E-PG neurons expressed the appropriate wildtype E-PG markers: R60D05-LexA, Toy, and Dac (Fig. 3C-D’’’, F). We conclude that Svp is required to restrict E-PG neuron production to the early T2NB window. We hypothesize that an early TTF may be required to generate the early born E-PG and P-EN neuron identities. We subsequently focus on the candidate TTF Castor.

**Figure 3.**
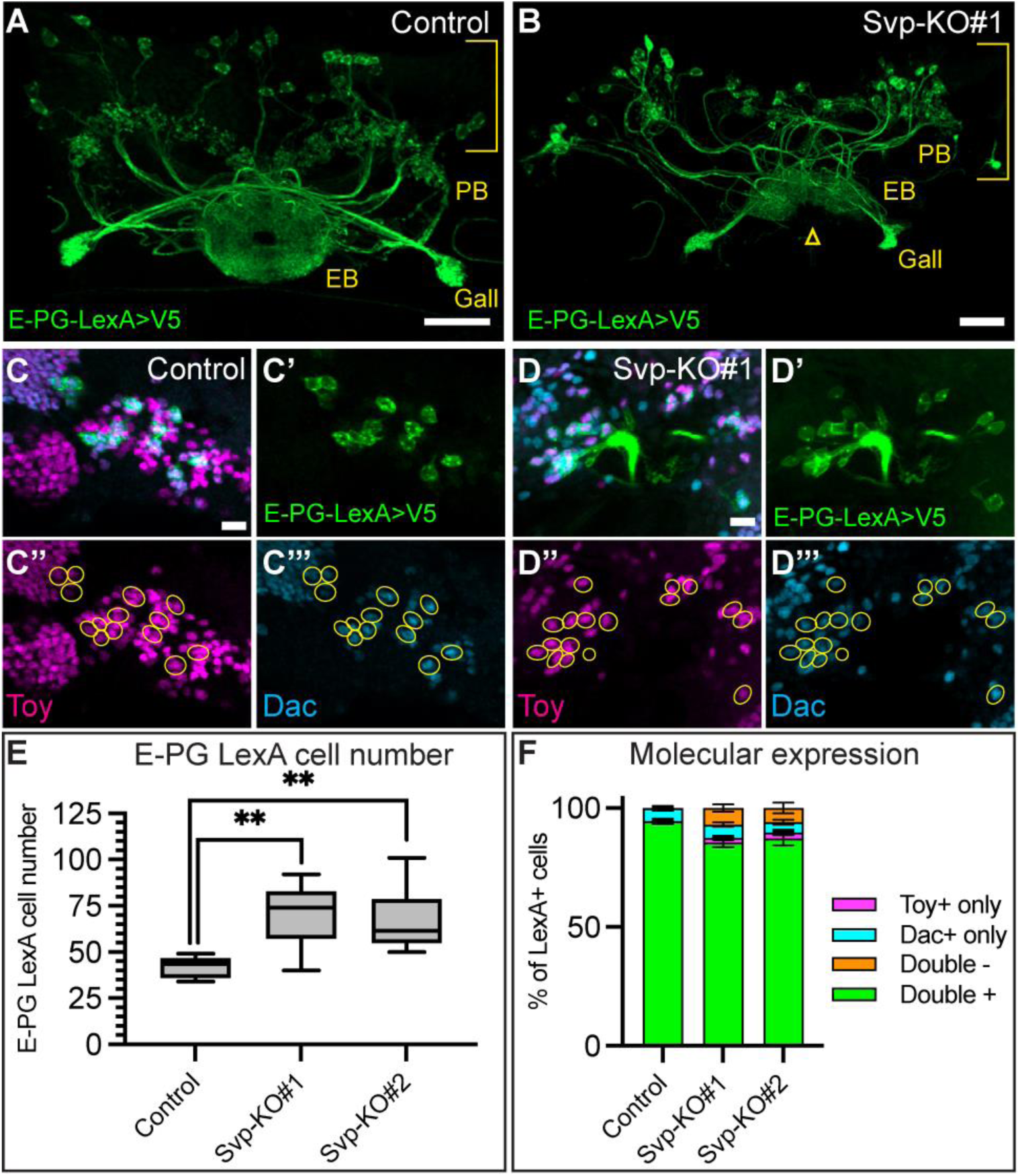
Seven-up is required to restrict E-PG production in early Type 2 neuroblast lineages. (A,B) Control (A) and Seven-up (Svp) knockout (B). There is an expansion of R60D05-LexA+ E-PG neurons in the Svp knockout. Brackets indicate cell body region; yellow text labels central complex neuropils: protocerebral bridge (PB), ellipsoid body (EB), and gall. Yellow arrowhead indicates morphological defect in the EB. (C-D) Control (C) and Svp knockout (D). Ectopic R60D05-LexA+ E-PG neurons maintain expression of molecular markers: R60D05-LexA, Toy, and Dac. (E) Quantification of LexA cell numbers from A,B. Box and whisker plots display the minimum and maximum range of the data with interquartile range. Control, *n*=16; Svp-KO#1, *n*=15; Svp-KO#2, *n*=14. *P*-values were determined using a one-way ANOVA, \*\**P*<0.001, followed by unpaired *t-*tests between the control and Svp-KOs: Control versus Svp-KO#1, \*\**P*<0.001; Control versus Svp-KO#2, \*\**P*<0.001. (F) Quantification of the molecular expression of E-PG neurons. Error bars show standard error of the mean. *n* the same as in E. In all images, LexA+ neurons driving membrane-bound V5 are in green. Neurons outlined in yellow in C-D’’’; Toy, magenta; Dac, cyan. Scale bars: 20 μm in A-B; 5 μm in C-D’’’.

### Castor expression is transient in the Type 2 larval neuroblast lineages

The early expression window of Castor in larval T2NBs makes it an ideal candidate as a TTF that specifies early born CX neuron identities. Before assaying neuron identity with Castor manipulations, we characterized the expression of Castor in the larval T2 lineage. We used Pnt-Gal4 to label the T2 lineage in larval stages (Zhu et al., 2011), Dpn to label NBs and INPs (NBs identified as ≥ 5 μm in diameter; INPs identified as < 5 μm) (Boone and Doe, 2008; Bowman et al., 2008), and Elav to label post-mitotic neurons (Robinow and White, 1991). We found that Castor was expressed in T2NBs only prior to 48 h ALH (Fig. 4A-D). INPs expressed Castor across 24-96 h ALH (Fig. 4E-H). T2 larval neurons expressed Castor starting at 48 h ALH with peak occurrence of Castor positive neurons at 72 h ALH before decreasing in occurrence at 96 h ALH (Fig. 4I-L). Castor expression was not maintained into the adult E-PG or P-EN neurons (Fig. 4M-O). We conclude that Castor is expressed in an early and narrow T2NB temporal window with transient expression across the larval T2 lineage and is not maintained in the adult P-EN and E-PG neurons (Fig. 4P). The Castor expression in T2NBs aligns with our birth dating of E-PG and P-EN neurons to the early temporal window (Fig. 1) (Dillon et al., 2024); thus, we focus on Castor in subsequent experiments.

**Figure 4.**
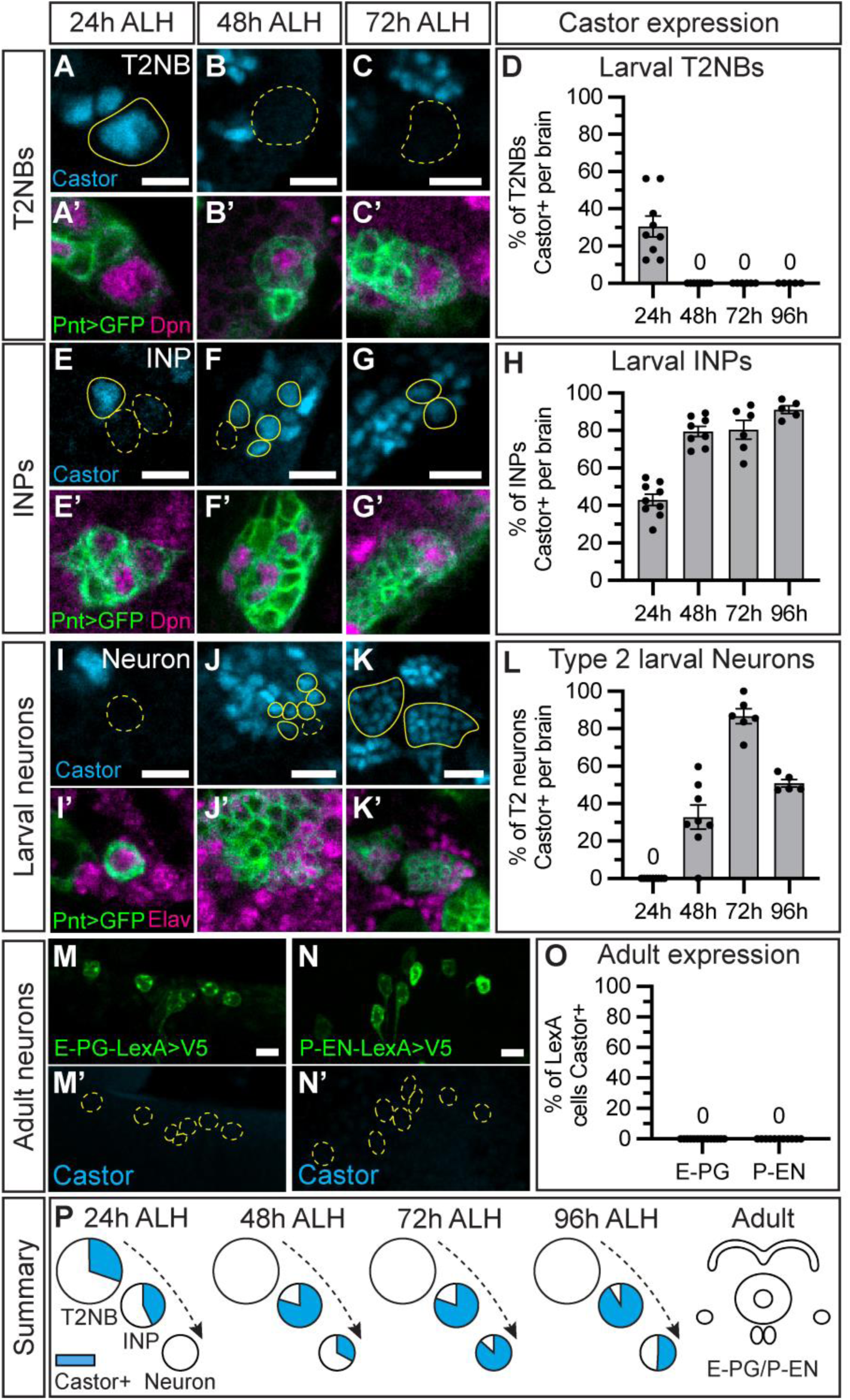
Castor expression is transient in Type 2 neuroblast lineages. (A-D) Castor expression in wild-type T2NBs at 24 h (A), 48 h (B), and 72 h ALH (C). Castor expression was seen at 24 h and at no later timepoints. (D) Quantification. Bar plot shows mean with error bars showing standard error of the mean (SEM). Each dot represents one brain. 24 h, *n*=9; 48 h, *n*=8; 72 h, *n*=7; 96 h, *n*=5. (E-H) Castor expression in wild-type INPs at 24 h (E), 48 h (F), and 72 h ALH (G). (H) Quantification. Bar plot shows mean with SEM. Each dot represents one brain. *n* the same as in D. (I-L) Castor expression in wild-type INPs at 24 h (I), 48 h (J), and 72 h ALH (K). (L) Quantification. Bar plot shows mean with SEM. Each dot represents one brain. *n* the same as in D. (M-O) Adult E-PG neurons (M-M’) and adult P-EN neurons do not express Castor (N-N’). LexA neurons driving membrane-bound V5, green. (O) Quantification. Bar plot. Each dot represents one brain. E-PG, *n*=14; P- EN, *n*=11. (P) Summary of Castor expression in the larval Type 2 neuroblast lineage and adult neurons shows transient expression across time and cell types. In all images, Pnt-Gal4>GFP+ cells in green, A-K’; LexA+ neurons in green M-N’; cells outlined in yellow; solid line indicates positive for Castor, dashed line indicates negative for Castor; Dpn or Elav, magenta; Castor, cyan. All scale bars: 5 μm.

### Generating Type 2 lineage specific Castor knockout and misexpression lines

To test if Castor is required and/or sufficient to specify CX neuron identities, we validated a Castor knockout (Castor-KO) line and two UAS-Castor misexpression (Castor ME) lines expressed with the Pnt-Gal4 driver. We generated a Castor-KO specific to the T2NB lineages. We used a CRISPR/Cas9 line (Port et al., 2020) with two Castor-specific sgRNAs to knockout *castor* in T2NBs. Similar to our previously reported Svp-KO lines (Dillon et al., 2024), we found that Castor-KO effectively knocks out *castor* in the majority of T2NBs by 24 h ALH, when the P-EN and E-PG neurons are first being born, with a few escaper T2NBs that show wild-type expression of Castor (Fig. S2A-C). We found two Castor misexpression lines (Castor-ME) (Kambadur et al., 1998) that ectopically extend Castor expression into 48 h ALH T2NBs (Fig. S2D-F). We conclude that these Castor-KO and Castor-ME lines are sufficient to test the role of Castor in specifying adult CX neuron identities.

To determine if Castor and Svp cross-regulate in the larval T2NB lineage, we used our Castor-KO and Svp-KO lines to assay the resulting expression of each temporal factor within the T2NB lineages. We found that Castor-KO did not disrupt the expression of Svp in 24 h ALH T2NBs (Fig. S3A-C). Additionally, we found that Svp- KO did not result in prolonged expression of Castor in T2NBs at 48 h ALH (Fig. S3D-F). These data support previous findings that Castor and Svp are not cross regulatory in the larval T2NBs (Ren et al., 2017; Syed et al., 2017).

### Castor is required to specify early born P-EN and E-PG adult neuron molecular identities

To determine if Castor is required as a TTF to specify early born neuron identities, we expressed Castor-KO in larval T2NB lineages and assayed P-EN and E-PG neuron identities in the adult. We found that a loss of Castor led to a loss of R12D09-LexA P-EN neurons (Fig. 5A-C). Additionally, Castor-KO led to a loss of the molecular markers Runt and Cut in the remaining P-EN neurons (Fig. 5D-F). We found similar phenotypes in E-PG neurons with Castor-KO resulting in a loss of R60D05-LexA E-PG neurons (Fig. 5G-I). Furthermore, Castor-KO showed a loss of the E-PG molecular markers Toy and Dac in the remaining E-PG neurons (Fig. 5J- L). We conclude that Castor is required for specifying the molecular identities of the adult P-EN and E-PG neurons.

**Figure 5.**
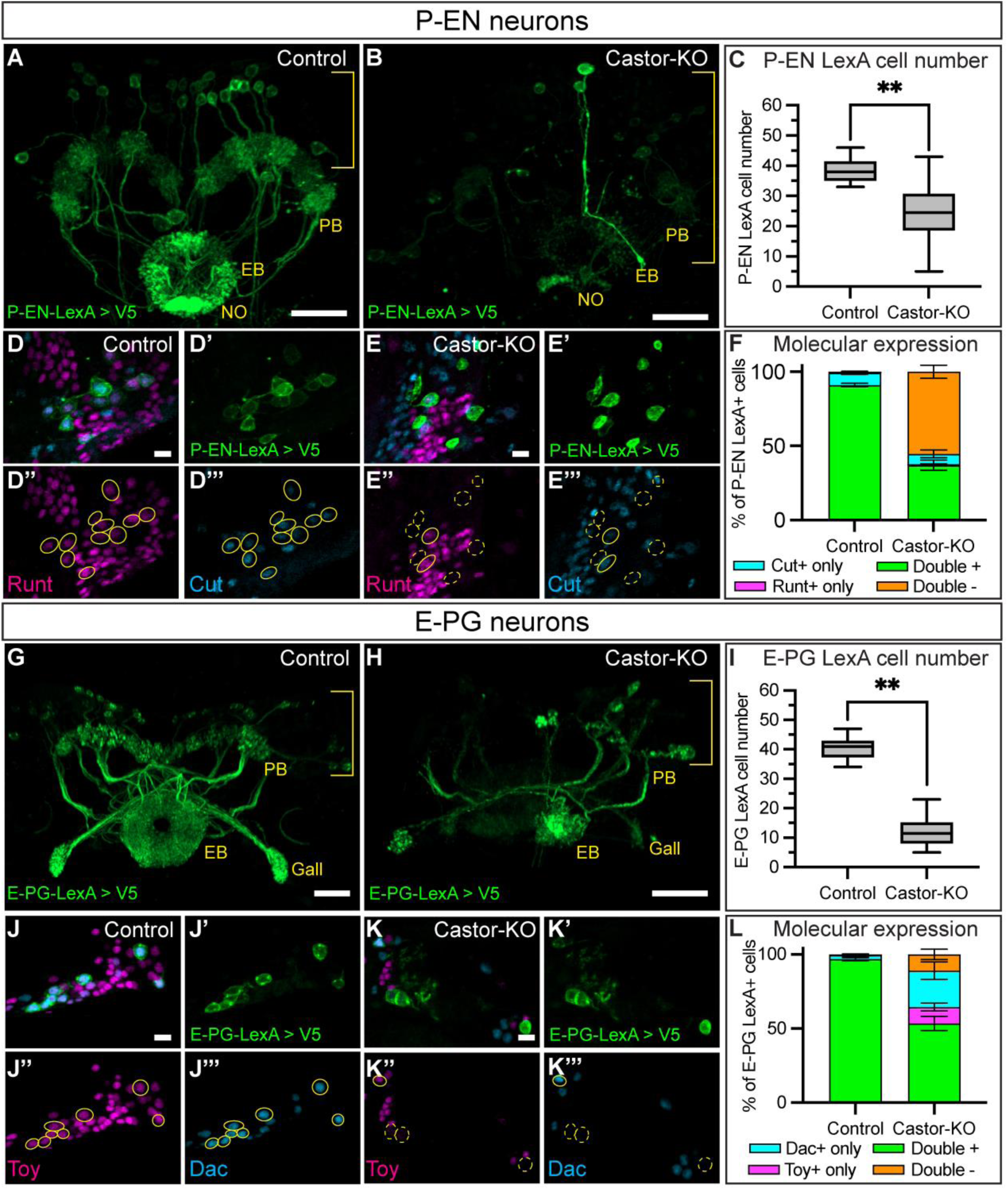
Castor is required to specify early born P-EN and E-PG adult neuron molecular identities. (A-C) Control (A) and Castor knockout (B) show a loss of R12D09-LexA+ P-EN neurons in the Castor knockout. Brackets indicate cell body region; yellow text labels central complex neuropils: protocerebral bridge (PB), ellipsoid body (EB), and noduli (NO). (C) Quantification. Box and whisker plots display the minimum and maximum range of the data with interquartile range. Control, *n*=17; Castor-KO, *n*=16. *P*-value was determined using an unpaired *t-*test, \*\**P*<0.001. (D-F) Control (D-D’’’) and Castor knockout (E-E’’’) show a loss of P-EN molecular markers with the loss of Castor. (F) Quantification. Bar plot shows mean with error bars showing standard error of the mean (SEM). *n* the same as in C. (G-I) Control (G) and Castor knockout (H). R60D05-LexA+ E-PG neurons are reduced in the Castor knockout. Brackets indicate cell body region; yellow text labels central complex neuropils: protocerebral bridge (PB), ellipsoid body (EB), and gall. (I) Quantification. Box and whisker plots display the minimum and maximum range of the data with interquartile range. Control, *n*=16; Castor-KO, *n*=18. *P*-value was determined using an unpaired *t-*test, \*\**P*<0.001. (J-L) Control (J-J’’’) and Castor knockout (K-K’’’) show a loss of E-PG molecular markers with the loss of Castor. (L) Quantification. Bar plot shows mean with SEM. *n* the same as in I. In all images, LexA+ neurons driving membrane- bound V5 are in green and outlined in yellow; solid line indicates double positive for markers, dashed line indicates not double positive for markers, Runt or Toy, magenta; Cut or Dac, cyan. Scale bars: 20 μm in A-B,G-H; 5 μm in D-E’’’, J-K’’’.

### Castor is sufficient to produce ectopic adult P-EN neurons but not E-PG neurons

To determine if Castor is sufficient to produce ectopic early born identities, we used our two Castor-ME lines in the T2NB lineage and assayed for adult P-EN and E-PG neurons. We found that Castor-ME led to an increase in R12D09-LexA P-EN neuron number (Fig. 6A-C). Additionally, these ectopic P-EN neurons maintained wild-type expression of the P-EN markers Runt and Cut (Fig. 6D-F). We found no noticeable phenotype with Castor-ME in R60D05-LexA E-PG neurons with no change in neuron number (Fig. 6G-I) or molecular expression (Fig. 6J-L). We conclude that Castor is sufficient to produce ectopic adult P-EN neurons but not E-PG neurons.

**Figure 6.**
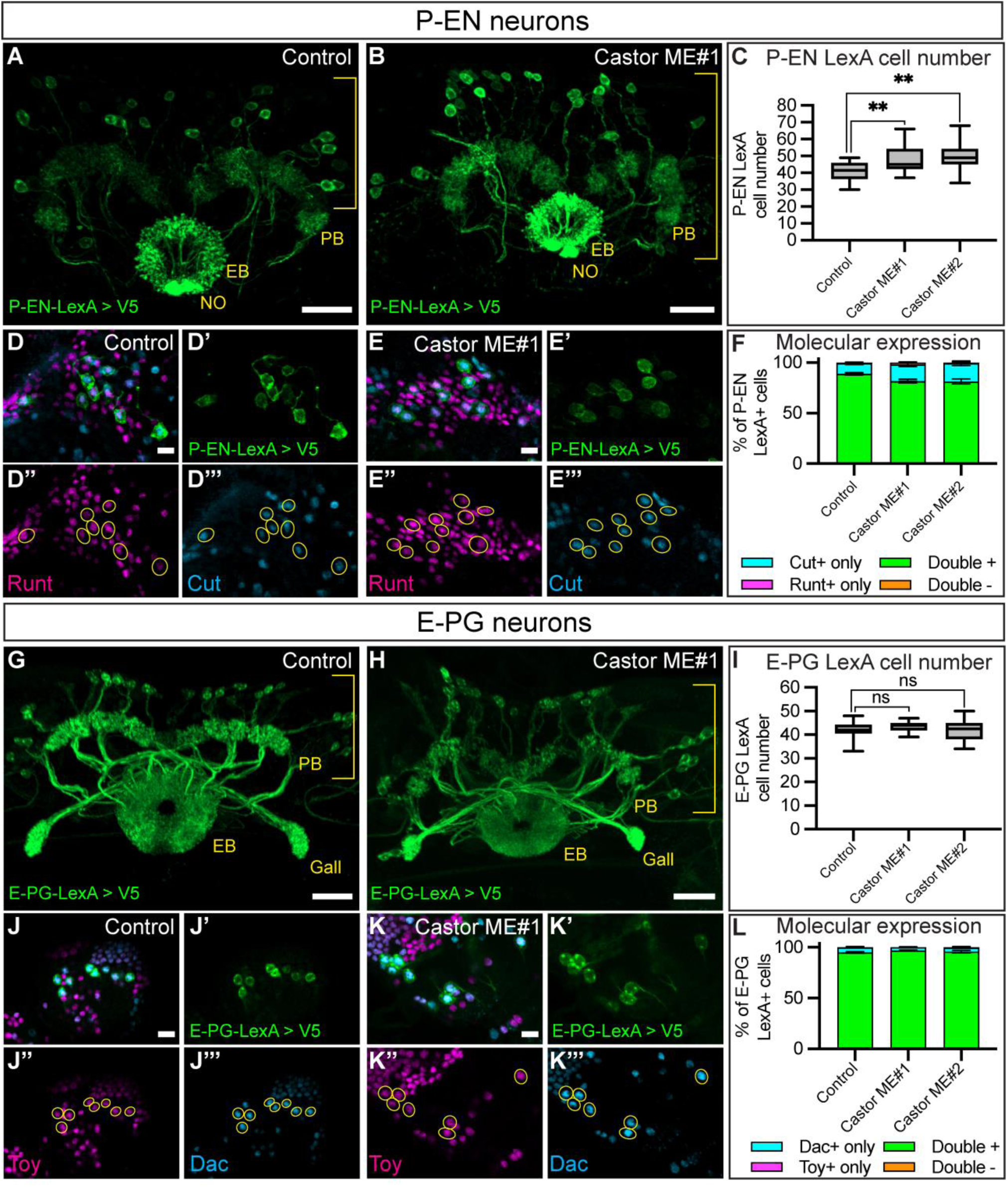
Castor is sufficient to produce ectopic adult P-EN neurons but not EPG neurons. (A-C) Control (A) and Castor misexpression (B) show a gain of R12D09-LexA+ P-EN neurons in the Castor misexpression (ME). Brackets indicate cell body region; yellow text labels central complex neuropils: protocerebral bridge (PB), ellipsoid body (EB), and noduli (NO). (C) Quantification. Box and whisker plots display the minimum and maximum range of the data with interquartile range. Control, *n*=16; Castor ME#1, *n*=16; Castor ME#2, *n*=18. *P*-values were determined using a one-way ANOVA, \*\**P*=0.005, followed by unpaired *t-*tests between the control and Castor misexpressions: Control versus Castor ME#1, \*\**P*=0.009; Control versus Castor ME#2, \*\**P*=0.002. (D-F) Control (D-D’’’) and Castor ME#1 (E-E’’’) shows a maintenance of P-EN molecular markers with the misexpression of Castor. (F) Quantification. Bar plot shows mean with standard error of the mean (SEM). *n* the same as in C. (G-I) Control (G) and Castor misexpression (H). R60D05-LexA+ E-PG neurons are unchanged in Castor ME. Brackets indicate cell body region; yellow text labels central complex neuropils: protocerebral bridge (PB), ellipsoid body (EB), and gall. (I) Quantification. Box and whisker plots display the minimum and maximum range of the data with interquartile range. Control, *n*=18; Castor ME#1, *n*=18; Castor-ME#2, *n*=18. *P*-value was determined using a one-way ANOVA, *P*=0.33. (J- L) Control (J-J’’’) and Castor-ME#1 (K-K’’’) show a no change in E-PG molecular markers with Castor-ME. (L) Quantification. Bar plot shows mean with SEM. *n* the same as in I. In all images, LexA+ neurons driving membrane-bound V5 are in green and outlined in yellow; Runt or Toy, magenta; Cut or Dac, cyan. Scale bars: 20 μm in A-B,G-H; 5 μm in D-E’’’, J-K’’’.

## Discussion

### Castor expression in larval Type 2 neuroblast lineages

We report a characterization of Castor across larval development and in most stages of the T2 lineage (NB, INP, and neurons). Previous work has only reported the T2NB expression window of Castor (Bayraktar and Doe, 2013; Ren et al., 2017; Syed et al., 2017). We confirmed previous findings that Castor expression in T2NBs is restricted to the early temporal window (Fig. 4A-D) (Bayraktar and Doe, 2013; Ren et al., 2017; Syed et al., 2017). We show that Castor is expressed outside of T2NBs in INPs and postmitotic larval neurons at later stages (Fig. 4E-H). Notably, the adult E-PG and P-EN neurons do not express Castor (Fig. 4M-O). These data are consistent with a TTF pattern of transient expression within a narrow temporal window in the parental NB, and inheritance of expression by progeny cells, as seen for embryonic TTFs (Cleary and Doe, 2006; Grosskortenhaus et al., 2006; Isshiki et al., 2001; Meng et al., 2019; Meng et al., 2020; Moris-Sanz et al., 2015; Novotny et al., 2002; Pearson and Doe, 2003; Seroka and Doe, 2019; Tran and Doe, 2008).

This indicates that Castor may play a role in specifying identities but is not required to maintain neuron identity. Determining Castor target genes in these transient stages remains an unanswered question. Future work is needed to identify the DNA-binding targets of Castor.

### Castor is a narrowly expressed early temporal transcription factor in larval Type 2 lineages

It remains an open question of how T2NBs generate diverse, birth order-dependent identities. The two hypotheses are that the lineage could use: (1) broad temporal gradients of protein expression similar to mushroom body neurons (Liu et al., 2015); or (2) a TTF cascade similar to the embryonic NB lineages (Cleary and Doe, 2006; Grosskortenhaus et al., 2006; Isshiki et al., 2001; Meng et al., 2019; Meng et al., 2020; Moris-Sanz et al., 2015; Novotny et al., 2002; Pearson and Doe, 2003; Seroka and Doe, 2019; Tran and Doe, 2008). Here, we show that Castor acts as an early TTF in the larval T2NB lineage for specifying early-born neuron identities (Fig. 5; Fig. 6). Additionally, previous work has shown that Castor and the switching factor Svp act upstream of these protein gradients (Ren et al., 2017). Recent work has also shown that these broadly expressed protein gradients are required for proper CX neuron specification in both T2NBs and INPs (Hamid et al., 2024; Munroe and Doe, 2023). These data are consistent with a model that the T2NB lineage is patterned by both briefly expressed TTFs and lengthy gradients of RNA-binding proteins (Fig. 1A). In the future, it will be important to determine whether other TTFs exist in T2NBs and how these temporal patterning genes are regulated.

One hallmark of TTFs have been their ability to form cross regulatory TTF cascades as seen in the embryonic lineages (Cleary and Doe, 2006; Grosskortenhaus et al., 2006; Isshiki et al., 2001; Meng et al., 2019; Meng et al., 2020; Moris-Sanz et al., 2015; Novotny et al., 2002; Pearson and Doe, 2003; Seroka and Doe, 2019; Tran and Doe, 2008). We, and others, report that Castor and Svp do not cross regulate in the larval T2NBs (Fig. S3) (Ren et al., 2017; Syed et al., 2017). This raises an open question of whether T2NBs use a TTF cascade or another mechanism of patterning to regulate temporal progression. Additional TTFs will need to be identified to test these hypotheses.

We show that Castor is required for the early born P-EN and E-PG adult neurons (Fig. 5); conversely, we show that Castor is sufficient to produce additional adult P- EN neurons, but not E-PG neurons (Fig. 6). These data, along with Castor’s early expression window, are consistent with Castor acting as an early TTF in the T2NB lineage. Interestingly, loss of Svp does not extend the Castor window (Fig. S3) (Ren et al., 2017; Syed et al., 2017) but does extend the production of early born neuron identities (Fig. 3) (Dillon et al., 2024). This could be due to a difference in the INP window in which P-EN and E-PG neurons are born (young vs old). Alternatively, additional early factors may be required for specifying fates and could explain the lack of Castor sufficiency to produce additional E-PG neurons. Future work should aim to determine additional molecular mechanisms for specifying these CX neuron identities.

### Columnar neurons born in the same Type 2 neuroblast window have distinct adult molecular identities

This work and previous work have shown that columnar neurons express distinct molecular markers that differentiate their identities (Dillon et al., 2024; Epiney et al., 2023; Sullivan et al., 2019). We show that E-PG and P-EN neurons are born within the same T2NB early window but have distinct molecular identities (Fig. 1C-E; Fig. 2) (Dillon et al., 2024). Interestingly, previous work has shown that columnar neurons born in the same INP windows share common markers: for example, young INP derived P-EN and P-FN neurons express Runt while old INP derived E-PG and PF-R neurons express Toy, even when these fates are born in different NB windows (Sullivan et al., 2019; Dillon et al., 2024). It remains an open question whether columnar neurons born in the same NB window share common molecular markers specified within the T2NB regardless of their INP birth window.

### Conserved role of Castor in vertebrate neurogenesis

Our work shows that Castor acts as a TTF in the early larval T2NBs. Previous work has shown Castor is a late TTF in embryonic NB lineages (Grosskortenhaus et al., 2006; Tran and Doe, 2008), although it has not been identified as a TTF in optic lobe NB lineages (El-Danaf et al., 2023). The mammalian ortholog of *castor*, *Casz1*, has been shown in mammalian models of the mouse retina and dorsal root ganglion to be temporally expressed in progenitors to control the generation of neuronal subtypes (Mattar et al., 2015; Mattar et al., 2018; Mattar et al., 2021; Monteiro et al., 2016). Furthermore, Casz1 has been shown in retinal progenitor cells to be regulated by Ikzf1 (Mattar et al., 2015), an ortholog of the *Drosophila* TTF Hunchback in embryonic NB lineages (Cleary and Doe, 2006; Grosskortenhaus et al., 2006; Isshiki et al., 2001; Meng et al., 2019; Meng et al., 2020; Moris-Sanz et al., 2015; Novotny et al., 2002; Pearson and Doe, 2003; Seroka and Doe, 2019; Tran and Doe, 2008). These findings suggest that Castor has a conserved role as a temporal patterning gene from fly to mammals (Frith et al., 2024; Liu et al., 2023; Sagner and Briscoe, 2019).

## Materials and Methods

### Animal Preparation

*Drosophila melanogaster* was used in all experiments. All flies were kept and maintained at 25°C unless stated otherwise. Stocks used can be found in Table. S1 and experimental genetic crosses in Table. S2.

#### EdU experiment

EDU (5-ethynyl-2′-deoxyuridine; Millipore-Sigma, 900584-50MG) was used to label proliferating cells starting at various sequential larval ages. Larvae were fed food containing 20 μg/ml EdU nonstop from the initial age feeding started until pupation. Larvae fed on EdU were raised at temperatures between 18°C and 21°C until adults hatched and were dissected.

#### Larval experiments

Embryos were collected on 3% agar apple juice caps with yeast paste for 4 hours and aged for the equivalent time after larval hatching (ALH). Hatched larvae were collected and dissected.

#### Adult experiments

Males and virgin females were introduced in standard yeast medium vials and flipped every two days. 2-5 day old adult flies were dissected for all experiments unless stated otherwise. All animals dissected were a mixture of male and female unless otherwise specified.

### Immunohistochemistry

Antibodies used with supporting notes can found in Table. S3.

#### Larval brain sample preparation

Larval brains were dissected in PBS and mounted on poly-L-lysine coated coverslips (Corning BioCoat 354085). Samples fixed for 23 minutes in 4% PFA in PBST. Samples were washed in PBST and blocked with 2% normal donkey serum (Jackson ImmunoResearch Laboratories, Inc. 017-000-121) in PBST. Samples incubated in a primary antibody mix diluted in PBST at room temperature for 4 h or at 4°C overnight. Primary antibodies were removed, and samples thoroughly washed with PBST. Samples were incubated in secondary antibodies at room temperature for 4 h or overnight at 4°C. Secondary antibodies were removed, and samples washed in PBST. Samples were dehydrated with an ethanol series of 30%, 50%, 75%, and 100% ethanol then incubated in xylene (Fisher Chemical X5-1) for 2x10 minutes. Samples were mounted onto slides with DPX (Sigma-Aldrich 06552) and cured for 3-4 days then stored at 4°C until imaged.

#### Adult brain sample preparation

Adult brains were prepared similar to larval brains with the exception of 38 minutes for fixation in 4% PFA and 2x12 minute xylene incubations.

#### EdU adult brain sample preparation

Adult brains from EdU-fed larvae were dissected in HL3.1 then fixed in 4% PFA for 30 min and incubated in block at 4°C overnight. Samples were incubated in primary and secondary mixes before Click-it-Reaction to label EdU. The Click-it-Reaction mix comprised PBS, Copper II sulfate (ThermoFisher, 033308.22), 555-Azide (ThermoFisher, A20012) in DSMO and ascorbic acid (Sigma-Aldrich, A4544-25G) for a 2h incubation. Samples were dehydrated and washed in xylene before DPX mounting as described above.

### Confocal Microscopy

Fixed preparations were imaged with a Zeiss LSM 900 laser scanning confocal (Carl Zeiss AG, Oberkochen, Germany) equipped with an Axio Imager.Z2 microscope. A 10x/0.3 EC Plan-Neofluar M27 or 40x/1.40 NA Oil Plan-Apochromat DIC M27 objective lens were used. Software program used was Zen 3.6 (blue edition) (Carl Zeiss AG, Oberkochen, Germany).

### Image processing and analysis

#### Figure preparation

Images in figures were prepared either in Imaris 10.0.1 or FIJI. Scale bars are given for a single slice in all single slice images and from all stacks within maximum intensity projections images. Pixel brightness was adjusted in images for clearer visualization; all adjustments were made uniformly over the entire image, and uniformly across wild-type samples and corresponding control and experimental samples. Adobe Illustrator 2024 (Adobe, Mountain View, CA) was used for figure formatting.

#### Statistical analyses

Statistics were computed using Prism 10 (GraphPad Software, Boston, MA). All statistical tests used are listed in the figure legends. *P*-values are reported in the figure legends. Plots display ns=not significant with *P*>0.05, \**P*<0.05, \*\**P*<0.01. Plots were generated using Prism with standard error of the mean bars shown and box and whisker plots display the minimum and maximum range of the data with interquartile range.

## Additional files

## Acknowledgements

We thank Laurina Manning and Jordan Munroe for assistance with EdU experiments and Sen-Lin Lai for advice on antibodies. We thank Tori Herman and fellow lab members Derek Epiney and Gonzalo Morales Chaya for comments on the manuscript. Antibodies obtained from the Developmental Studies Hybridoma Bank, created by the NICHD of the NIH and maintained at the University of Iowa, Department of Biology, Iowa City, IA were used in this study. Stocks obtained from the Bloomington Drosophila Stock Center (NIH P40OD018537) and Vienna Drosophila Resource Center were used in this study.

## Author contributions

Conceptualization: N.R.D.; Methodology: N.R.D.; Validation: N.R.D.; Formal analysis: N.R.D.; Investigation: N.R.D.; Resources: N.R.D.; Data curation: N.R.D.; Writing - original draft: N.R.D.; Writing - review & editing: N.R.D., C.Q.D.; Visualization: N.R.D.; Supervision: C.Q.D.

## Data availability

All relevant data can be found within the article and its supplementary information.

---

**Supplemental Fig 1.**
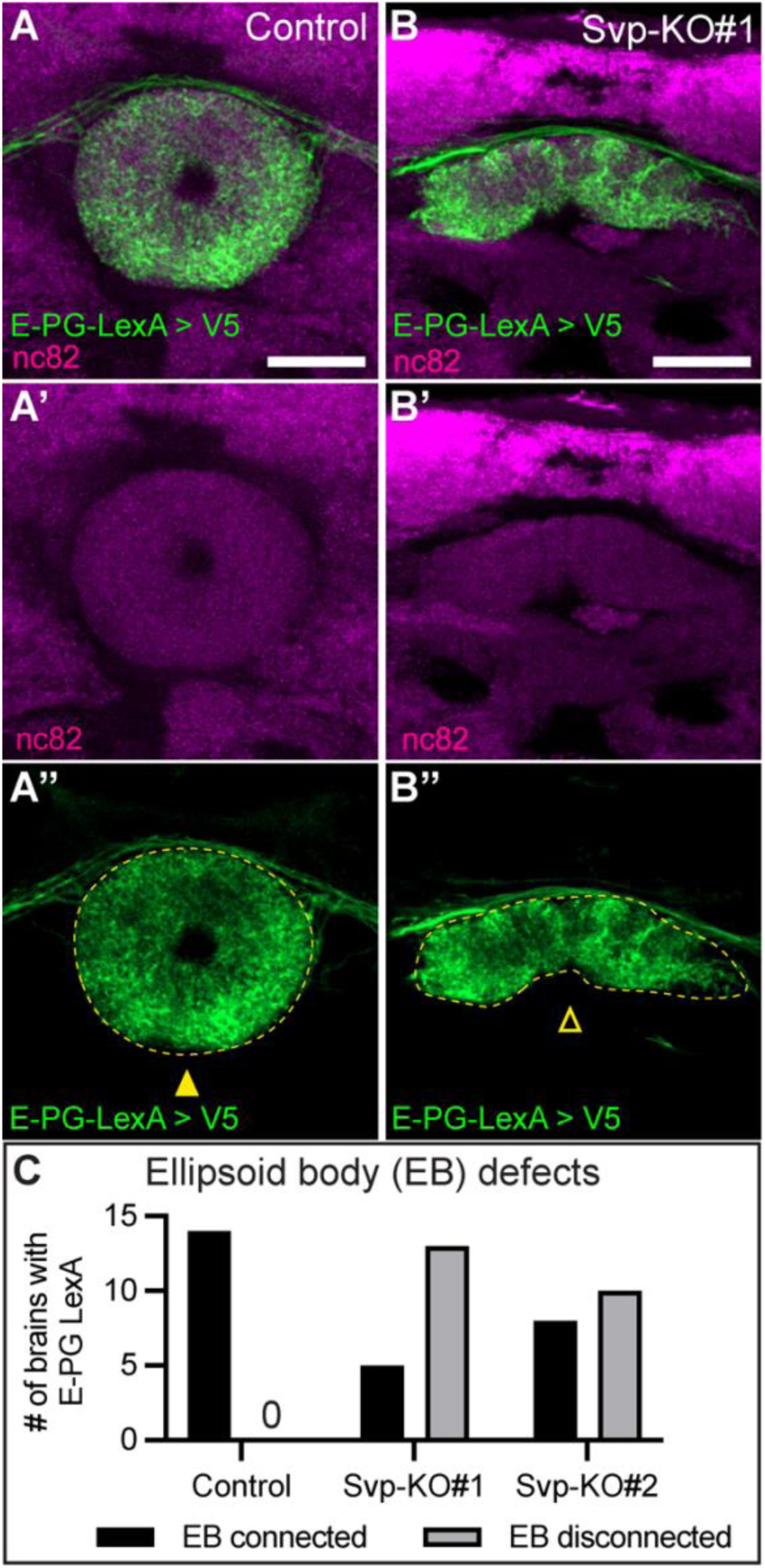
Loss of Seven-up in Type 2 neuroblasts leads to altered E-PG ellipsoid body morphology. (A-B) Control (A) and Seven-up (Svp) knockout (B). There is altered morphology of R60D05-LexA+ E-PG neurons in the Svp knockout with a disconnected ellipsoid body (EB). (C) Quantification. Control, *n* = 14; Svp-KO#1, *n* = 18; Svp-KO#2, *n* = 18. *P*-value determined by Chi-square analysis ** *P*<0.001. In all images, LexA+ neurons driving V5 expression are in green, neuropil nc82 in magenta, and EB outlined in yellow dashed line. Closed arrowhead indicates a connected EB; open arrowhead indicates a disconnected EB. Scale bars: 20 μm.

**Supplemental Fig 2.**
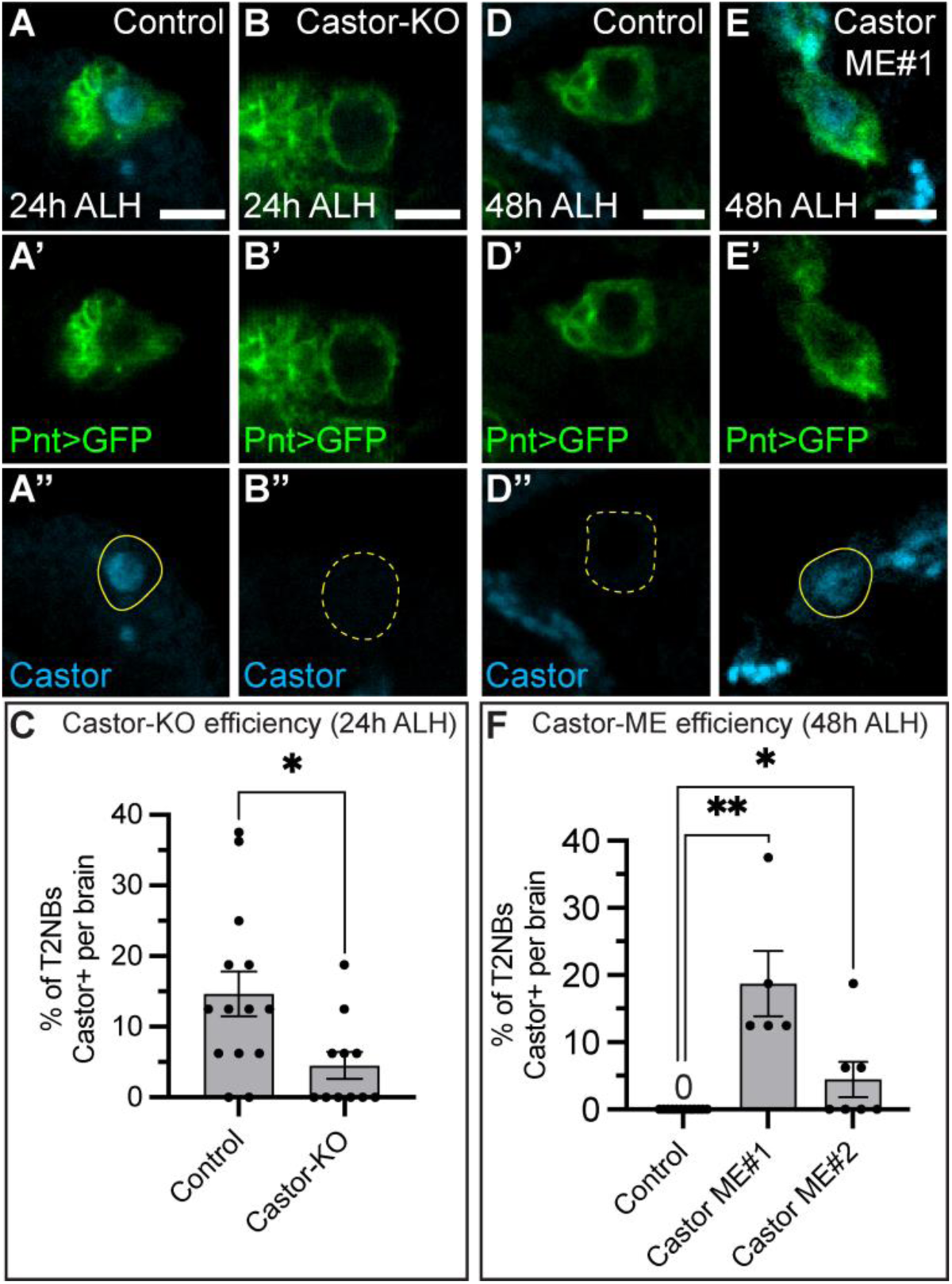
Generating Type 2 lineage specific Castor knockout and misexpression lines. (A-C) Control (A) and Castor-KO (B) show a loss of Castor in Type 2 neuroblasts (T2NB) in the Castor-KO at 24h after larval hatching (ALH). (C) Quantification. Bar plot shows mean with standard error of the mean (SEM). Each dot represents one brain. Control, *n*=14; Castor-KO, *n*=11. *P*-value was determined using an unpaired *t-* test, \**P*=0.017. (D-F) Control (A) and Castor misexpression (ME) (B) show ectopic expression of Castor in Type 2 neuroblasts (T2NB) in the Castor-ME at 48h after larval hatching (ALH). (F) Quantification. Bar plot shows mean with SEM. Each dot represents one brain. Control, *n*=12; Castor 2nd, *n*=5; Castor 3rd, *n*=7. *P*-values were determined using a one-way ANOVA, \*\**P*<0.001, followed by unpaired *t-*tests between the control and Castor-MEs: Control versus Castor 2nd, \*\**P*<0.001; Control versus Castor 3rd, \**P*=0.036. In all images, Pnt-Gal4 driving GFP in green and T2NBs outlined in yellow; solid line indicates positive for Castor, dashed line indicates negative for Castor; Castor, cyan. Scale bars: 5 μm.

**Supplemental Fig 3.**
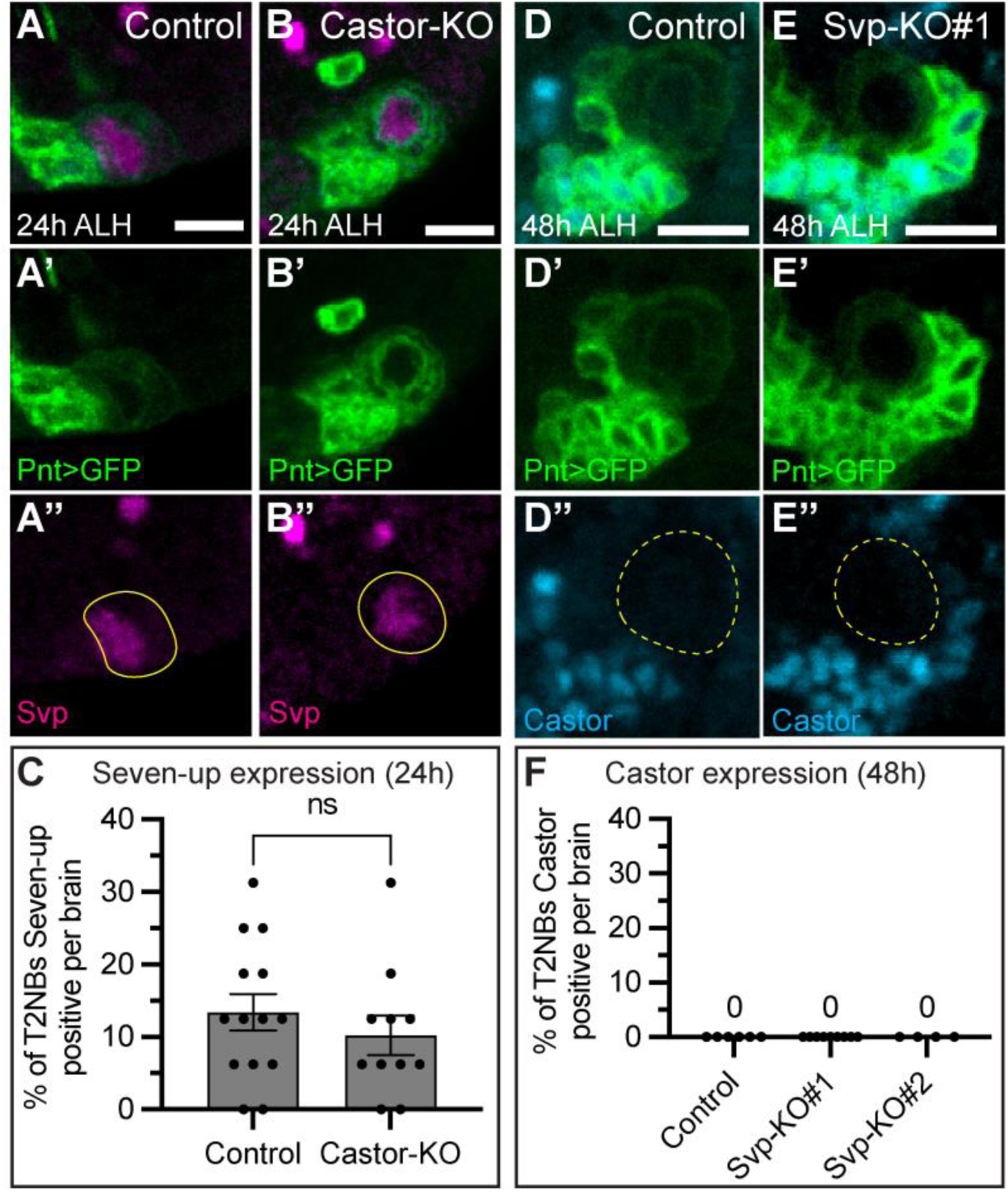
Castor and Seven-up do not cross regulate in Type 2 neuroblasts. (A-C) Control (A) and Castor-KO (B) shows no change in Seven-up (Svp) expression in Type 2 neuroblasts (T2NB) in the Castor-KO at 24 h ALH. (C) Quantification. Bar plot shows mean with standard error of the mean. Each dot represents one brain. Control, *n*=14; Castor-KO, *n*=11. *P*-value was determined using an unpaired *t-*test, \**P*=0.40. (D-E) Control (D) and Svp-KO#1 (E) shows no extended expression of Castor in Type 2 neuroblasts (T2NB) in the Svp-KO at 48 h ALH. (F) Quantification. Bar plot shows mean with no error bars due to no differential values. Each dot represents one brain. Control, *n*=6; Svp-KO#1, *n*=9; Svp-KO#2, *n*=4. *P*-value was not determined due to no differential values reported between conditions. In all images, Pnt-Gal4 driving GFP in green and T2NBs outlined in yellow; solid line indicates positive for transcription factor of interest, dashed line indicates negative for transcription factor of interest; Svp, magenta; Castor, Cyan. Scale bars: 5 μm.

**Supplementary table 1.** Transgenes and *Drosophila melanogaster* stock lines used.

| **Genotype** | **Source and identifier** | **Additional information** |
| --- | --- | --- |
| *R12D09-LexA* | BDSC #54419 | Expressed in P-EN neurons |
| *R60D05-LexA* | BDSC # 52867 | Expressed in E-PG neurons |
| *13xLexAop-myr::GFP* | BDSC #32210 | Expresses membrane bound GFP under LexAop control |
| *10XUAS-IVS-myr::GFP* | BDSC #32198 | Expresses membrane bound GFP under UAS control |
| *Pointed-Gal4* | PMID: 22143802<br>14-94 | Expressed in Type 2 lineage starting in the neuroblast |
| *10xUAS-IVS-myr::smGdP::HA, 13xLexAop2-IVS-myr::smGdP::V5* | BDSC #64092 | Expresses HA membrane tag under UAS control, V5 membrane tag under LexAop control |
| *hsFLP; ;UAS-Cas9.P2* | BDSC #58986 | Expresses Cas9 under UAS control |
| *hsFLP; UAS-sgRNA::svp ;* | VDRC #341527 | Expresses two short guide RNAs against Svp under UAS control; Svp-KO#1 |
| *hsFLP; UAS-sgRNA::svp ;* | VDRC #341390 | Expresses two short guide RNAs against Svp under UAS control; Svp-KO#2 |
| *hsFLP; UAS-sgRNA::castor ;* | VDRC #341386 | Expresses two short guide RNAs against Castor under UAS control |
| *; UAS-LacZ ;* | BDSC #8529 | Expresses LacZ under UAS control |
| *; UAS-Castor ;* | Odenwald, W. | Expresses Castor under UAS control |
| *; ; UAS-Castor* | Odenwald, W. | Expresses Castor under UAS control |

**Supplementary table 2.** Genetic crosses for each experiment.

| **Figures** | **Summary** | **Genetic cross** |
| --- | --- | --- |
| Figure 1C-E | Labels adult E-PG neurons for EdU birth dating | Females containing *13xLexAop-myr::GFP* were crossed to males containing *R60D05-LexA* |
| Figure 2A-C<br>Figure 4M-O | Labels adult E-PG (i) and P-EN (ii) neurons for molecular markers | Self-cross of females and males containing (i) *10xUAS-IVS-myr::smGdP::HA*, *13xLexAop2-IVS-myr::smGdP::V5*; *R60D05-LexA*; *Pointed-Gal4* or (ii) *10xUAS-IVS-myr::smGdP::HA*, *13xLexAop2-IVS-myr::smGdP::V5*; *R12D09-LexA*; *Pointed-Gal4* |
| Figure 3A-F<br>Figure S1A-C | Labels adult E-PG neurons and drives UAS constructs in the Type 2 progenitors for: Cas9.P2 control (i) Svp-KO#1 (ii) Svp-KO#2 (iii) | Females containing *10xUAS-IVS-myr::smGdP::HA*, *13xLexAop2-IVS-myr::smGdP::V5*; *R60D05-LexA*; *Pointed-Gal4* were crossed to males containing either (i) ; ; *UAS-Cas9.P2*, (ii) ; *UAS-sgRNA::svp* (VDRC #341527) ; *UAS-Cas9.P2* , or (iii) ; *UAS-sgRNA::svp* (VDRC #341390) ; *UAS-Cas9.P2* |
| Figure 4A-L | Labels larval Type 2 lineage | Self-cross of females and males containing *10XUAS-IVS-myr::GFP* ; *Pointed-Gal4* |
| Figure 5A-F | Labels adult P-EN neurons and drives UAS constructs in the Type 2 progenitors for: Cas9.P2 control (i) Castor-KO#1 (ii) | Females containing *10xUAS-IVS-myr::smGdP::HA*, *13xLexAop2-IVS-myr::smGdP::V5*; *R12D09-LexA*; *Pointed-Gal4* were crossed to males containing either (i) ; ; *UAS-Cas9.P2*, (ii) ; *UAS-sgRNA::castor*; *UAS-Cas9.P2* |
| Figure 5G-L | Labels adult E-PG neurons and drives UAS constructs in the Type 2 progenitors for: Cas9.P2 control (i) Castor-KO#1 (ii) | Females containing *10xUAS-IVS-myr::smGdP::HA*, *13xLexAop2-IVS-myr::smGdP::V5*; *R60D05-LexA*; *Pointed-Gal4* were crossed to males containing either (i) ; ; *UAS-Cas9.P2*, (ii) ; *UAS-sgRNA::castor*; *UAS-Cas9.P2* |
| Figure 6A-F | Labels adult P-EN neurons and drives UAS constructs in the Type 2 progenitors for: LacZ control (i) Castor ME#1 (ii) Castor ME#2 (iii) | Females containing *10xUAS-IVS-myr::smGdP::HA*, *13xLexAop2-IVS-myr::smGdP::V5*; *R12D09-LexA*; *Pointed-Gal4* were crossed to males containing either (i) ; *UAS-LacZ* ; (ii) ; *UAS-Castor* ; (iii) ; ; *UAS-Castor* |
| Figure 6G-L | Labels adult E-PG neurons and drives UAS constructs in the Type 2 progenitors for:<br>LacZ control (i)<br>Castor ME#1 (ii)<br>Castor ME#2 (iii) | Females containing *10xUAS-IVS-myr::smGdP::HA*, *13xLexAop2-IVS-myr::smGdP::V5*; *R60D05-LexA*; *Pointed-Gal4* were crossed to males containing either (i) ; *UAS-LacZ* ; (ii) ; *UAS-Castor* ; (iii) ; ; *UAS-Castor* |
| Figure S2A-F | Labels larval Type 2 lineage and drives UAS constructs in the Type 2 progenitors for:<br>Cas9.P2 control (i)<br>Castor-KO#1 (ii)<br>LacZ control (iii)<br>Castor ME#1 (iv)<br>Castor ME#2 (v) | Females containing *10XUAS-IVS-myr::GFP* ; *Pointed-Gal4* were crossed to males containing either (i) ; ; *UAS-Cas9.P2*, (ii) ; *UAS-sgRNA::castor*, *UAS-Cas9.P2* (iii) ; *UAS-LacZ* ; (iv) ; *UAS-Castor* ; (v) ; ; *UAS-Castor* |
| Figure S3A-C | Labels larval Type 2 lineage and drives UAS constructs in the Type 2 progenitors for:<br>Cas9.P2 control (i)<br>Castor-KO (ii) | Females containing *10XUAS-IVS-myr::GFP* ; *Pointed-Gal4* were crossed to males containing either (i) ; ; *UAS-Cas9.P2*, (ii) ; *UAS-sgRNA::castor*, *UAS-Cas9.P2* |
| Figure S3D-F | Labels larval Type 2 lineage and drives UAS constructs in the Type 2 progenitors for:<br>Cas9.P2 control (i)<br>Svp-KO#1 (ii)<br>Svp-KO#2 (iii) | Females containing *10XUAS-IVS-myr::GFP* ; *Pointed-Gal4* were crossed to males containing either (i) ; ; *UAS-Cas9.P2*, (ii) ; *UAS-sgRNA::svp* (VDRC #341527) ; *UAS-Cas9.P2* , or (iii) ; *UAS-sgRNA::svp* (VDRC #341390) ; *UAS-Cas9.P2* |

**Supplementary table 3.** Antibodies used.

| **Antibody** | **Source and identifier** | **Additional information** |
| --- | --- | --- |
| Chicken anti-GFP | Aves: 1020 | (1:1000) |
| Rabbit anti-Castor | PMID:1418995; W. Odenwald | (1:1000) |
| Mouse anti-Seven-up 6F7 | DSHB: Hiromi, Y. / Hondo, T.<br>/ Kanda, H. | (1:4) |
| Rat anti-Deadpan | Abcam: 11D1 BC7.1B | (1:20) |
| Guinea pig anti-Runt | Claude Desplan lab (NYU) | (1:1000) |
| Guinea pig anti-Toy | Genscripts<br>U1806FG070_27/DC0126 | (1:1000) |
| Mouse anti-Cut 2B10 | DSHB: Rubin, G.M. | (1:10) |
| Mouse anti-Dac mAbdac1-1 | DSHB: Rubin, G.M. | (1:50) |
| Mouse anti-Elav 9F8A9 | DSHB: Rubin, G.M. | (1:50) |
| Mouse anti-V5 tag | ThermoFisher: R960-25<br>(previously Invitrogen 46-0705) | (1:1000) |
| Rabbit anti-V5 tag | Cell signaling: 13202S | (1:1000) |
| Mouse anti-nc82 | DSHB: Buchner, E. | (1:100) |
| Alexa Fluor® 488 AffiniPure Donkey Anti-Chicken IgY (IgG) (H+L) | Jackson ImmunoResearch, West Grove, PA: 703-545-155 | (1:400) |
| Rhodamine Red™-X (RRX) AffiniPure Donkey Anti-Rat IgG (H+L) | Jackson ImmunoResearch, West Grove, PA: 712-295-153 | (1:400) |
| Alexa Fluor® 647 AffiniPure Donkey Anti-Mouse IgG (H+L) | Jackson ImmunoResearch, West Grove, PA: 715-605-151 | (1:400) |
| Rhodamine Red™-X (RRX) AffiniPure Donkey Anti-Guinea pig IgG (H+L) | Jackson ImmunoResearch, West Grove, PA: 706-295-148 | (1:400) |
| Alexa Fluor® 488 AffiniPure Donkey Anti-Mouse IgG (H+L) | Jackson ImmunoResearch, West Grove, PA: 715-295-151 | (1:400) |
| Alexa Fluor® 488 AffiniPure Donkey Anti-Rabbit IgG (H+L) | Jackson ImmunoResearch,<br>West Grove, PA: 711-295-152 | (1:400) |
| Rhodamine Red™-X (RRX) AffiniPure Donkey Anti-Mouse IgG (H+L) | Jackson ImmunoResearch,<br>West Grove, PA: 715-295-151 | (1:400) |
| Alexa Fluor® 405 AffiniPure Donkey Anti- Mouse IgG (H+L) | Jackson ImmunoResearch,<br>West Grove, PA: 715-475-150 | (1:200) |

**Supplemental Table 4.** Raw data for experiments done. Each tab shows the raw data for the indicated figure or portion of a figure.

